# Cross-species real time detection of trends in pupil size fluctuation

**DOI:** 10.1101/2024.02.12.579393

**Authors:** Sharif I. Kronemer, Victoria E. Gobo, Catherine R. Walsh, Joshua B. Teves, Diana C. Burk, Somayeh Shahsavarani, Javier Gonzalez-Castillo, Peter A. Bandettini

## Abstract

Pupillometry is a popular method because pupil size is easily measured, sensitive to central neural activity, and associated with behavior, cognition, emotion, and perception. Currently, there is no method for online monitoring phases of pupil size fluctuation. We introduce *rtPupilPhase* – an open source software that automatically detects trends in pupil size in real time, enabling novel implementations of real time pupillometry towards achieving numerous research and translational goals. We validated the performance of rtPupilPhase on human, rodent, and monkey pupil data and propose future applications of real time pupillometry.

## Main Text

Pupillometry – the measure of pupil size – has grown in popularity with evidence that pupil size is an easily measured marker of neurophysiological activity. For instance, neuromodulatory networks, including the locus coeruleus-noradrenergic and basal forebrain-cholinergic systems, are linked to changes in pupil size (Figure 1A)^1,2^. These brain networks are also involved in widespread regulation of cortical and subcortical regions^3,4^. Therefore, pupillary dynamics can reflect both large-scale brain activity and behavioral, cognitive, emotive, and perceptual states that emerge by central neural mechanisms. For example, pupil size predicts states of locomotion, arousal, and conscious awareness^5–7^.

**Figure 1.**
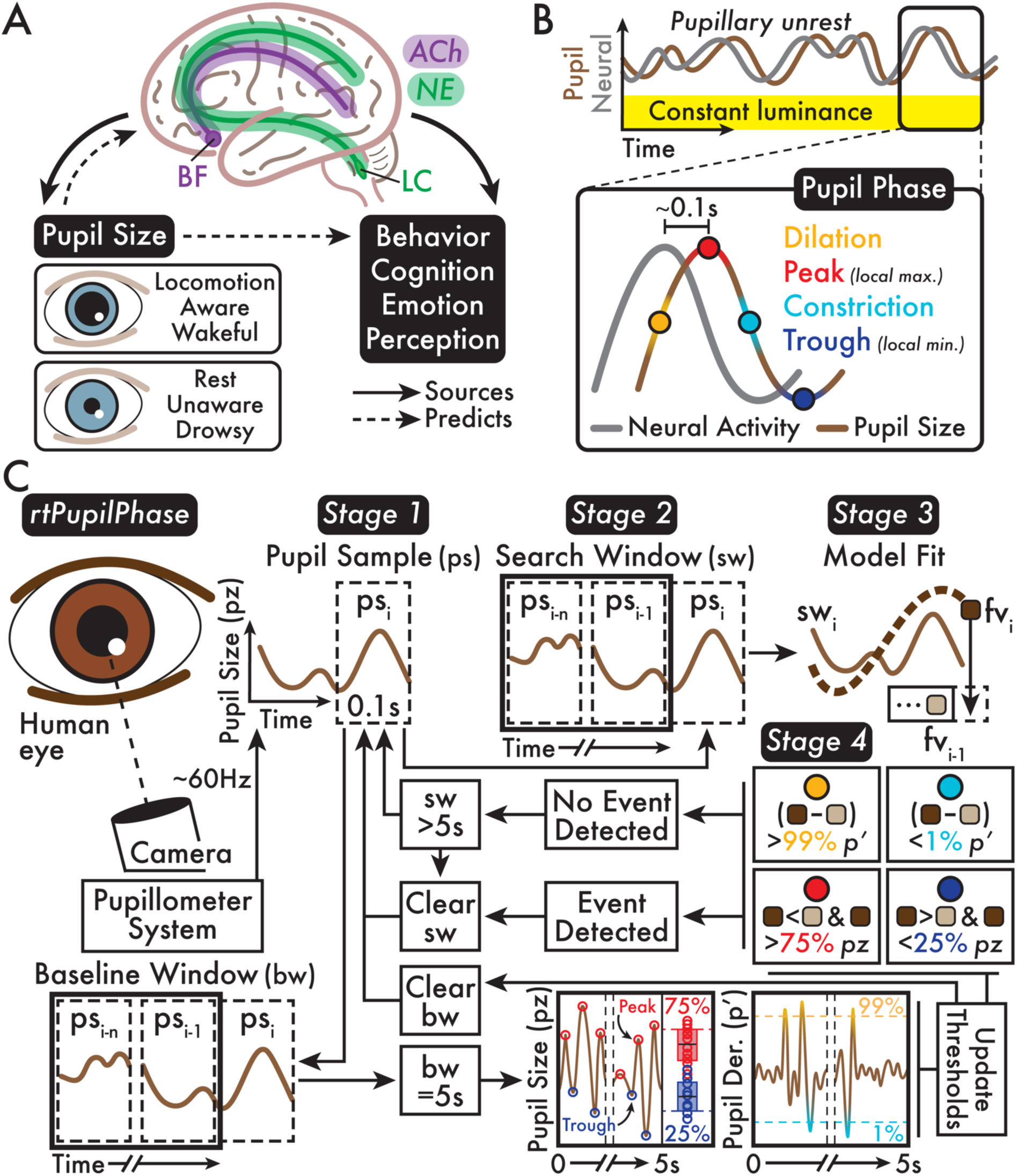
Neural sources and phases of pupil size fluctuation and the rtPupilPhase method schematic summary. **(A)** Major neuromodulatory networks, including the basal forebrain (BF) cholinergic system (ACh) and the locus coeruleus (LC) noradrenergic system (NE), broadly modulates subcortical and cortical activity. In addition, spontaneous and evoked ACh and NE neural activity are linked with pupil size. Therefore, pupil size can predict both large-scale brain activity and behavioral, cognitive, emotive, and perceptual states. For example, in humans, dilated pupils indicate locomotion, awareness, and wakefulness, while constricted pupils indicate rest, unawareness, and drowsiness. **(B)** *Pupillary unrest* is the spontaneous fluctuation of pupil size independent of environmental luminance. These trends in pupil size or *pupil phase* are a lagging (∼0.1 seconds [s]) indicator of neural activity. Although depicted as a sine wave, the light-independent trend of pupillary unrest is a heterogeneous, chaotic signal. Pupil size fluctuations are summarized in four main phases: (1) *dilation*, (2) *peak* or local maximum (max), (3) *constriction*, and (4) *trough* or local minimum (min). **(C)** The *rtPupilPhase* method is summarized in four stages. *Stage 1:* Live streamed pupil data (∼60 Hz sampling rate; see *Human Pupillometry Acquisition* Methods section) is gathered into *pupil samples* (ps) of 0.1-second intervals. *Stage 2:* The ps is added to a *search window* (sw) and *baseline window* (bw). *Stage 3:* The sw data is modeled with a quadratic fit. The final pupil size value (fv) from the fitted sw model is recorded. *Stage 4:* A pupil phase event is determined by comparing the fv from the current and previous sw and pupil size and pupil size derivative thresholds that are updated approximately every 5 s according to analysis of the bw. Dilation: fv_i_ minus fv_i-1_ is *greater than* the dilation pupil derivative threshold (p’); peak: fv_i_ is *less than* fv_i-1_ *and* fv_i_ is *greater than* the peak pupil size (pz) threshold; constriction: fv_i_ minus fv_i-1_ is *less than* the constriction pupil derivative threshold (p’); trough: fv_i_ is *greater than* fv_i-1_ *and* fv_i_ is *less than* the trough pupil size threshold. The sw is reset whenever an event was detected or the sw exceeds 5 s. See the *Real Time Detection of Pupil Phase Events* Methods section for full details.

The central neural underpinnings of pupil size explain *pupillary unrest*: fluctuations in pupil size under constant environmental luminance (Figure 1B)^8,9^. Pupillary unrest has a dominant frequency near 0.5 Hz, although stimulus evoked pupillary fluctuations (e.g., a flickering light) can induce pupil size changes up to ∼3.5 Hz^10–12^. While pupil size fluctuations do not conform to a perfect sine wave (e.g., ^13^), pupil size trends are summarized in four main phases: (1) *dilation*, (2) *peak* (i.e., local maximum), (3) *constriction*, and (4) *trough* (i.e., local minimum; Figure 1B). Previous research finds that these pupil phases are a lagging (∼0.1 seconds) indicator of central neural activity (Figure 1B)^1,4,6,14,15^.

The sensitivity of pupil phase to brain activity has encouraged the widespread recording of pupil size in myriad experimental contexts and across species. Pupillometry is commonly recorded passively and analyzed post hoc (e.g., correlating pupil size with behavioral performance and neuroimaging signals). Retrospective pupillometry analyses restrict experimental design and the interpretation of results. Alternatively, online or real time pupillometry – detecting trends in pupil size at or near the moment of their occurrence – would enable new task types (e.g., closed-loop paradigms) and novel applications (e.g., self-regulation of arousal state via pupil size feedback)^16^. Despite the potential advantages, there is currently no published procedure for the real time detection of pupil phase.

To address this deficit, we introduce *rtPupilPhase* – an open source software that automatically detects dilation, peak, constriction, and trough pupil phase events in real time. rtPupilPhase also identifies pupil phase-independent or *random* events that we used to evaluate the performance of rtPupilPhase. In future applications, random events could also serve as a phase-independent control condition. Monitoring pupil phase in real time is challenging, in part, because pupil size change is sluggish, undergoes baseline shifts, and the regular occurrence of behaviors that occlude the pupil (e.g., blinking and whisking in rodents). rtPupilPhase mounts these challenges and achieves the real time detection of pupil phase events in four main stages (Figure 1C; see *Real Time Detection of Pupil Phase Events* Methods section). In summary, rtPupilPhase aggregates intervals of live-streamed pupil size data, models these data intervals, and records detected pupil phase events determined by recurrently updated, individual-specific pupil size and pupil size derivative (i.e., pupil size samples x_1_-x_2_, x_2_-x_3_, etc.) thresholds.

We evaluated rtPupilPhase using data from healthy, adult human participants (N = 8; see *Participants* Methods section) who completed a passive fixation task integrated with rtPupilPhase under constant luminance and simultaneous head-fixed, pupillometry recording (see *Fixation Task* and *Human Pupillometry Acquisition* Methods sections). rtPupilPhase detected hundreds of pupil phase events in real time per participant (Supplementary Figure 1). The median pupil phase inter-event duration was 0.067 seconds or a ∼15 Hz detection rate (Figure 3B; Supplementary Figure 2A; see *Pupil Phase Event Detection Temporal Performance* Methods section). In fact, the majority (67.91%) of pupil phase inter-event durations were less than 0.1 seconds (Figure 3B). Therefore, the temporal performance of rtPupilPhase tested with the current parameters (Table 1) supports its use to monitor both spontaneous and evoked pupillary dynamics that are estimated to fluctuate at frequencies between ∼0.5 and 3.5 Hz (e.g., ^10,12^).

**Table 1.** rtPupilPhase and sim-rtPupilPhase parameters. For the monkey dataset, the sim-rtPupilPhase baseline window duration and inter-event interval were adjusted to 0.5 and 1.5 seconds, respectively. The percentile (%) thresholds are applied on pupil size and pupil derivative data from the baseline window and used to detect pupil phase events. *Human sim-rtPupilPhase data were down sampled from 1000 to 60 Hz to match the approximate online sampling rate while testing rtPupilPhase on humans. The variable online sampling rate of the human pupillometry recordings (see *Human Pupillometry Acquisition* Methods section) meant there was variability in the duration of the pupil sample, max search window, and baseline window. See the *Real Time Detection of Pupil Phase Events* and *Human*, *Mouse*, and *Monkey Simulated Real Time Detection of Pupil Phase Events* Methods sections for details.

|  | *Sampling Rate* | *Pupil Sample Duration* | *Max Search Window Duration* | *Baseline Window Duration* | *Inter-Event Interval* | *Pupil Phase Percentile Thresholds* |
| --- | --- | --- | --- | --- | --- | --- |
| Human (N = 8)<br>*rtPupilPhase* | ~60 Hz | 0.1 seconds | 5 seconds | 5 seconds | 3 seconds | *Peak: 75%*<br>*Trough: 25%*<br>*Dilation: 99%*<br>*Constriction: 1%* |
| Human (N = 8)<br>*sim-rtPupilPhase* | 1000 Hz* | 0.1 seconds | 5 seconds | 5 seconds | 3 seconds | *Peak: 75%*<br>*Trough: 25%*<br>*Dilation: 99%*<br>*Constriction: 1%* |
| Mouse (N = 5)<br>*sim-rtPupilPhase* | ~20 Hz | 0.1 seconds | 5 seconds | 5 seconds | 1 seconds | *Peak: 75%*<br>*Trough: 25%*<br>*Dilation: 99%*<br>*Constriction: 1%* |
| Monkey (N = 2)<br>*sim-rtPupilPhase* | 1000 Hz | 0.1 seconds | 5 seconds | 0.5 seconds | 1.5 seconds | *Peak: 75%*<br>*Trough: 25%*<br>*Dilation: 99%*<br>*Constriction: 1%* |

We extracted epochs of pupil size data centered on these pupil phase events (see *Human Pupil Size, Blink, and Saccade Fraction Epoch Extraction* Methods section). The resulting pupil size epoch timecourses demonstrated that rtPupilPhase reliably located pupil phases with high-temporal precision at the event (Supplementary Figure 2), participant (Supplementary Figure 3A), and group levels (Figure 2A; Supplementary Figure 4A). In addition, the pupil size timecourses significantly differed (cluster-based permutation testing, *p* < 0.05; see *Cluster-Based Permutation Testing* Methods section) between pupil phase and random events, particularly in the interval immediately preceding the pupil phase event (Supplementary Figure 4A).

**Figure 2.**
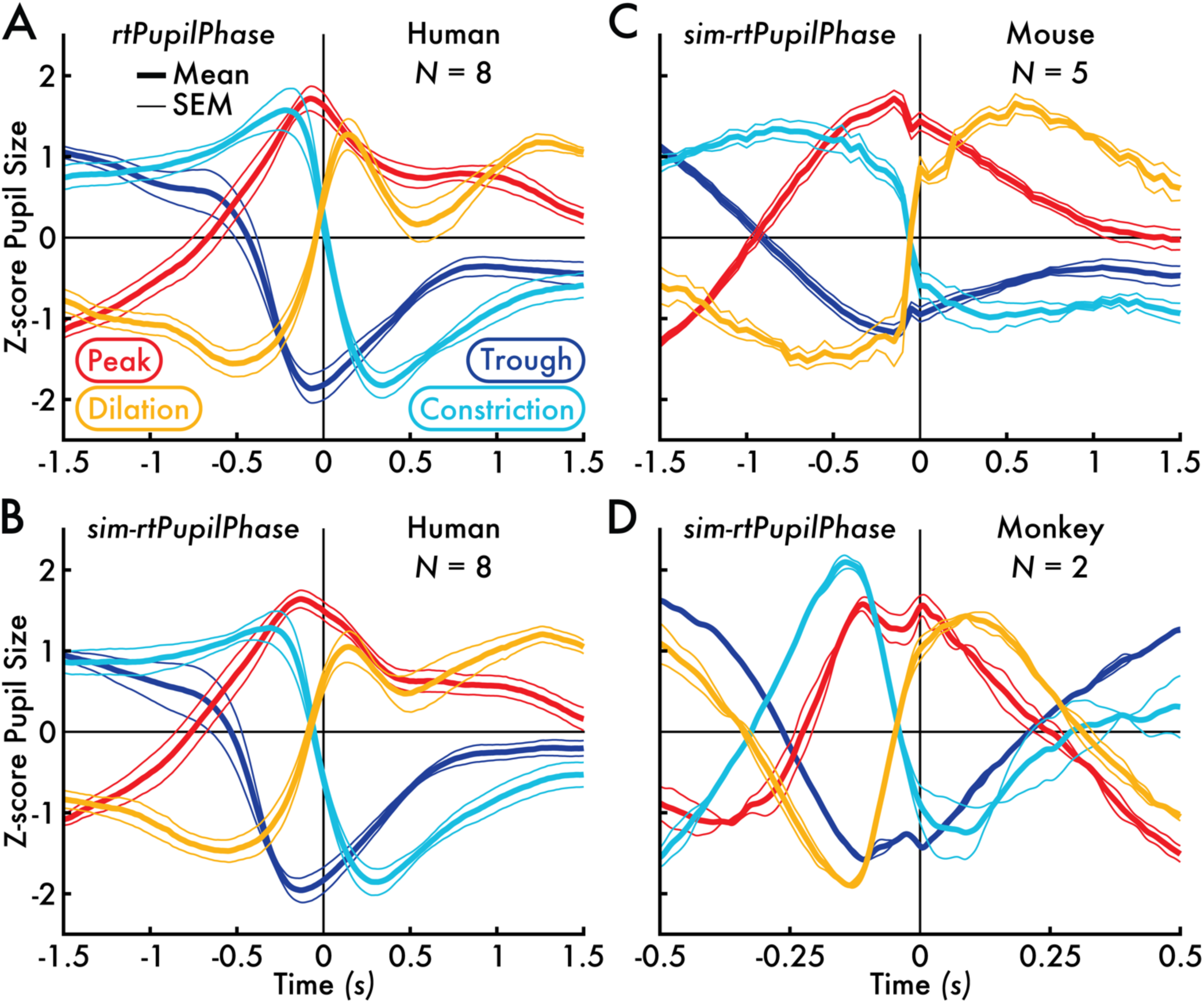
Human, mouse, and monkey mean z-score pupil size timecourses centered on pupil phase events detected with rtPupilPhase and sim-rtPupilPhase. **(A)** The rtPupilPhase human (N = 8) mean z-score pupil size timecourses. **(B)** The sim-rtPupilPhase human (N = 8) mean z-score pupil size timecourses. **(C)** The sim-rtPupilPhase mouse (N = 5) mean z-score pupil size timecourses. **(D)** The sim-rtPupilPhase monkey (N = 2) mean z-score pupil size timecourses. The pupil phase event time is 0 seconds (s). The group mean pupil phase event pupil size timecourse is shown in the thicker trace, bounded by thinner traces that depicts the standard error of the mean (SEM). The trace color corresponds with the pupil phase event type.

To assess the accuracy of rtPupilPhase, we compared the real time detected pupil phase events against the *true* pupil phase events, as determined by post hoc evaluation of the pupil data (see *Pupil Phase Event Detection Accuracy and Sensitivity Performance* Methods section). rtPupilPhase achieved an average pupil phase event-level accuracy of 88.16 (dilation), 79.26 (peak), 86.90 (constriction), and 73.37 percent (trough), all statistically greater than the random event accuracy of approximately 50 percent (*p* < 0.0005; Figure 3A). In addition, rtPupilPhase achieved high sensitivity. The on-target accuracy (i.e., comparing *detected* pupil phase events with the corresponding *true* pupil phase events; e.g., detected dilation versus true dilation; Figure 3A) was significantly greater (*p* < 0.002) than the off-target sensitivity (i.e., comparing *detected* pupil phase events with *true off*-target pupil phase events; e.g., detected dilation versus true peak) across all pupil phases (Supplementary Figure 5). In addition, the off-target sensitivity decreased with distance from the on-target pupil phase according to the pupil phase order. For example, the on-target detected dilation accuracy was 88.16 percent, followed by the off-target sensitivities of 61.27, 11.84, and 34.30 percent for peak, constriction, and trough events, respectively (Supplementary Figure 5A). Altogether, these results support that rtPupilPhase correctly identifies pupil phases near the moment of their true occurrence in real time.

**Figure 3.**
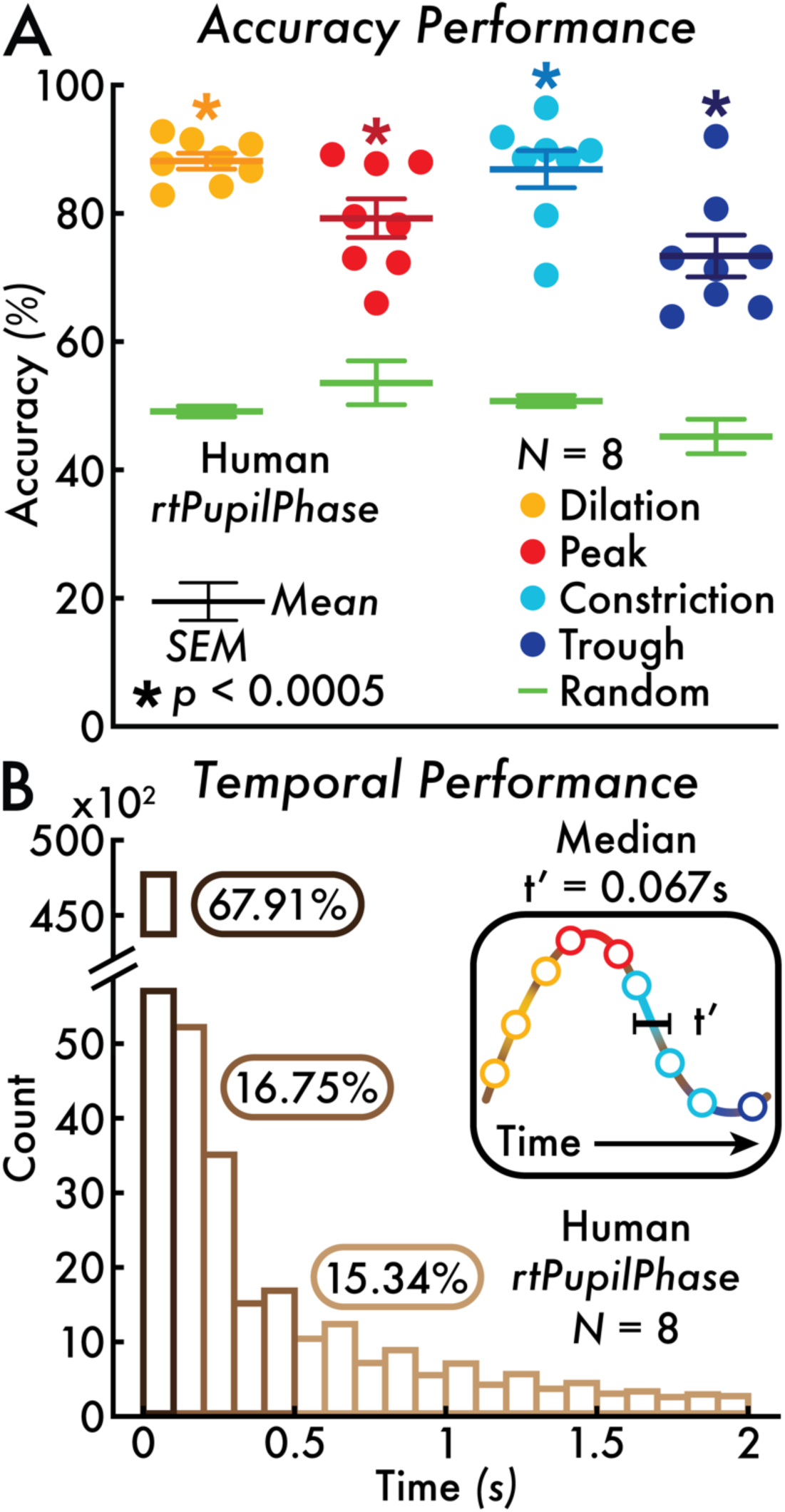
rtPupilPhase detected pupil phase event accuracy and temporal performance. **(A)** The pupil phase accuracy percentage for each human participant (each circle corresponds with a participant; N = 8) is shown for the rtPupilPhase real time detected pupil phase events versus the true pupil phase events. The random event accuracy relative to the true pupil phase event is shown in green (individual participant accuracy is not shown). The random accuracy percentage served as the baseline to statistically test against the pupil phase event accuracy (participant-level paired *t*-test; see *Pupil Phase Event Detection Accuracy and Sensitivity Performance* Methods section). All pupil phase event accuracies were statistically greater than the random event accuracy (Holm-Bonferroni corrected, *p* < 0.05; * *p* < 0.0005). The mean and standard error of the mean (SEM) are depicted with error bars. **(B)** The distribution of pupil phase inter-event durations (t’; see inset and *Pupil Phase Event Detection Temporal Performance* Methods section) across all detected pupil phase events up to 2 seconds (s) in binned increments of 0.1 s. The median t’ was 0.067 s. The majority of t’ values (67.91%) where less than 0.1 s; 16.75% between 0.1 and 0.5 s; and 15.34% greater than 0.5 s.

We also explored the rate of blinks and saccades preceding the real time detected pupil phase events, as eye movements are known to affect pupil size^17^. Blink and saccade occurrence epochs centered on the pupil phase events detected with rtPupilPhase were extracted. The resulting epoch timecourses averaged within event type and across participants revealed increases in blink and saccade fraction preceding the event time, each event type with different periods of peak change (Supplementary Figure 6C, E). To determine if the performance of rtPupilPhase was explained by blinks and saccades, we removed all pupil size epochs with blinks *or* saccades in the 0.5 seconds preceding the real time detected pupil phase events. Nonetheless, the pupil size timecourse trends among the remaining event epochs were unchanged across pupil phases (Supplementary Figure 6A). This result supports that rtPupilPhase detects pupil size trends independent of eye movements.

Next, we aimed to evaluate if rtPupilPhase could be used on pupillometry from non-human animals. As a proof of concept, we developed *sim-rtPupilPhase* – a software that *simulates* rtPupilPhase offline using previously collected pupil data (see *Human Simulated Real Time Detection of Pupil Phase Events* Methods section). We verified that sim-rtPupilPhase replicates the performance of rtPupilPhase using the previously described human pupil data. Although, achieving an exact duplication of the rtPupilPhase result with sim-rtPupilPhase was not possible due to the variable online sampling rate of the human pupillometry recordings (see *Human Pupillometry Acquisition* Methods section). Still, a similar number of pupil phase events were detected with sim-rtPupilPhase as with rtPupilPhase (Supplementary Figure 1B). Crucially, the extracted epochs centered on the pupil phase events detected with sim-rtPupilPhase reproduced the trends and statistics across pupil size, blink, and saccade fraction timecourses from events detected with rtPupilPhase (Figure 2A versus B; Supplementary Figure 3A versus B; Supplementary Figure 4A versus B; Supplementary Figure 6A versus B, C versus D, and E versus F).

After validating on humans, we implemented sim-rtPupilPhase to estimate the performance of rtPupilPhase on previously acquired pupil data from mice (N = 5; ^18^) and rhesus macaques (N = 2; data newly published here). As with the human dataset, sim-rtPupilPhase detected hundreds of pupil phase events for each non-human animal (Supplementary Figure 1C, D). Likewise, the trend of the pupil size epoch timecourses centered on these events was consistent with the human event pupil size timecourses detected with rtPupilPhase on the individual animal (Supplementary Figure 3C, D) and group levels (Figure 2C, D; Supplementary Figure 4C, D). These non-human animal results were achieved with minimal adjustment to sim-rtPupilPhase and rtPupilPhase parameters despite major differences among the datasets, including pupil physiology, experimental setup, task demands, and recording system (Table 1; see *Mouse* and *Monkey Simulated Real Time Detection of Pupil Phase Events* Methods sections).

In summary, event, participant, and group-level analyses support that rtPupilPhase was accurate and sensitive for predicting pupil phase in real time and across species. Future users of rtPupilPhase are encouraged to adapt and expand this method to improve performance and achieve unique applications. Assisting in future optimization, the fixation task (see *Fixation Task* Methods section) can be called from the command line interface with options to change the rtPupilPhase parameters (Table 1). Likewise, sim-rtPupilPhase can be used as a tool for testing the rtPupilPhase parameters without collecting additional data.

There are numerous potential applications of rtPupilPhase. A promising future implementation is the development of closed-loop paradigms (e.g., the real time detection of pupil phases triggers task events). This approach allows for targeted testing and possible causal interpretations for the interaction among pupil size and phase, neurophysiology, and behavior. For example, a study found that increasing pupil size at the time of a stimulus corresponded with improved perceptual performance, suggesting that the trend in pupil size is relevant to predict perceptual state^19^. Future experiments could extend on this finding by testing if pupil phase predicts perceptual-sensitivity by presenting near-threshold stimuli timed with target pupil phase events. Concurrent neuroimaging recording could help deduce the mechanism that may explain any changes in behavioral performance (e.g., perception rate and reaction time) linked with pupil phase. To encourage this kind of experimentation, we supplied a sample task script integrated with rtPupilPhase that could trigger an experimenter-defined task event (e.g., stimulus presentation) concurrent with the real time detection of pupil phases.

An alternative application of rtPupilPhase is as a tool for pupil biofeedback. A recent study found participants can self-regulate absolute pupil size with implications on brain activity^16^. Building on this study, using rtPupilPhase, participants could receive real time input of their pupil phase to regulate their pupillary fluctuations. Modifying pupil phase states may alter arousal, salience, attention, and perceptual brain network activity with corresponding consequences on behavior, cognition, emotion, and perception. Likewise, a future translational application of real time pupillometry biofeedback is as a method to regulate brain arousal network state as a possible treatment approach for people with impaired arousal level (e.g., sleep disorders and disorders of consciousness).

When considering future implementations of rtPupilPhase, there are several challenges to consider. First, users must be mindful of the pupil foreshortening error – changes in recorded pupil size due to head and eye movements independent of physiological state. In addition, luminance stimulation (e.g., very dark or bright environments) may impact the dynamic range of spontaneous or evoked pupillary fluctuations with consequences on detecting pupil phase events. Correspondingly, without additional performance testing, we recommend the use of rtPupilPhase with head stabilized pupillometry and in constant, neutral luminance environments. Also, the tested human and mouse pupil data where acquire during rest. Users who intend to implement rtPupilPhase during a task should confirm optimal pupil phase event detection for task-evoked pupillary fluctuations. As a proof of concept, the tested monkey pupil data were gathered during a task, suggesting rtPupilPhase can be robust in both rest and task contexts.

Finally, pupillary unrest is a continuous, chaotic signal that does not conform to a perfect sine wave^13^. Therefore, there is uncertainty for the ground truth definitions that determine the exact boundary among the pupil phases. Here, true dilation and constrictions phases was decided by the pupil size gradient, while true peak and trough phases were defined as all samples within 0.25 seconds of a local maxima or minima that exceeded a threshold prominence (see *Pupil Phase Event Detection Accuracy and Sensitivity Performance* Methods section). This approach for defining pupil phases resulted in overlapping event classifications (i.e., a single pupil size sample may belong to multiple pupil phase events). Likewise, we found that the detected pupil phase events corresponded with more than one event type (e.g., 61.27% of detected dilation events were within 0.25 seconds of a true peak event; Supplementary Figure 5). Alternative ground truth definitions of pupil phase events would necessarily change the rtPupilPhase performance metrics. Correspondingly, rtPupilPhase users must carefully study their pupil data and monitor the behavior of rtPupilPhase with diverse performance metrics.

In conclusion, real time pupillometry is a nascent approach with exciting applications. rtPupilPhase is a novel, open source method for monitoring trends in pupil size linked with behavioral, cognitive, emotive, and perceptual states across species. Several tools are provided that allow for seamless implementation and adaptation of rtPupilPhase to achieve optimized performance for unique application goals. An immediate use of rtPupilPhase includes developing closed-loop and biofeedback paradigms where the real time detection of pupil phase events triggers experimental stimuli. These types of paradigms are powerful to efficiently target pupil phases of interest and deduce casual relationships. Thus, rtPupilPhase introduces novel opportunities to study and implement the pupil-neurophysiology link towards achieving broad experimental and translational goals.

## Methods

### Participants

Healthy, adult humans (N = 8; mean age = 25.1 years; age standard deviation [SD] = 4.01 years; males = 1; mean education = 17.0 years; education SD = 1.9 years) were recruited from the local Bethesda, Maryland, USA community. The Institutional Review Board of the National Institute of Mental Health approved the recruitment and consent protocols. Inclusion criteria included: (1) being between 18 and 65 years old at the time of experimentation, (2) a healthy physical examine within a year of the study session, and (3) able to give informed consent. Exclusion criteria included: (1) no previous nor current histories of neurologic or psychiatric disorders, (2) poor vision that cannot be corrected, and (3) head injuries (e.g., loss of consciousness for >30 minutes and three or more concussive injuries). Each participant underwent a health exam completed by a nurse practitioner prior to their study session.

### Fixation Task

The fixation task was coded in Python and run with PsychoPy (v2022.2.4; Open Science Tools Ltd.). The task consisted of a single display: a central fixation point (a plus sign; 0.51 x 0.51 degrees) on a solid gray screen. Participants were instructed to maintain central fixation throughout the task. Participants completed five 10-minute fixation task blocks with a ∼2-minute break between each block. Integration of rtPupilPhase with the fixation task enabled the covert detection of pupil phase and random events (see *Real Time Detection of Pupil Phase Events* section) and accommodates future development of alternative experimental designs involving rtPupilPhase.

### Human Pupillometry Acquisition

Human, head-fixed pupillometry and eye tracking was acquired with a desktop mounted EyeLink 1000 Plus (recorded eye: right eye; offline sampling rate = 1000 Hz; Version 5.50; SR Research, Inc.). The online sampling rate of the pupil size data was variable (mode sampling rate ≈ 60 Hz or 17 milliseconds between samples). During pupillometry acquisition, participants were instructed to place their head in a chin and forehead rest system (SR Research Head Support; SR Research, Inc.) to maintain a fixed head position and a constant distance between the participant and pupillometer.

### Human Testing Equipment and Facility

Human testing was conducted in a single 1.5-hour study session in a windowless, temperature-controlled behavioral testing room. The lighting in the testing room was set to a consistent level for all participants and maintained throughout the entire study session. The fixation task was administered on a behavioral laptop (MacBook Pro; 13-inch; 2560 x 1600 pixels, 2019; Mac OS Catalina v10.15.7; Apple, Inc.)^20^. The EyeLink 1000 Plus software ran on a Dell desktop computer (Dell OptiPlex XE2; Dell, Inc.). The behavioral laptop and the fixation task were programmed to communicate with the EyeLink desktop via an Ethernet connection. Participants viewed the fixation task on a VIEWPixx display monitor (1920 x 1200 pixels; VPixx Technologies, Inc.). The behavioral laptop monitor was mirrored to the display monitor via a DVI cable. Participants were positioned approximately 56 centimeters from the center of the display monitor, measured from the nasion.

### Mouse Pupil Data

The mouse pupil data were acquired from a previously published experiment (https://zenodo.org/records/7968402)^18^. In summary, *in vivo* wide-field optical mapping, pupil size, locomotion, and whisking data were recorded from five adult male Thy1-jRGECO1a transgenic mice over multiple recording sessions. Throughout these sessions, the mice were head-fixed and free to exhibit spontaneous behaviors such as walking and running on a wheel without any experimental stimuli. The mouse pupil size was recorded with a BFS-U3-16S2M-CS USB 3.1 Blackfly S Monochrome Camera and sampled at ∼20 Hz. Using the tracking software DeepLabCut^21^, pupil size was monitored by identifying eight circumferential points, then fitting a circle to these points to estimate pupil diameter. For the current experiment, eight 10-minute recording sessions were selected for each mouse. Any session in which significant portions of the pupil data were missing was not selected for inclusion in the mouse dataset.

### Monkey Pupil Data

The monkey pupil data consisted of two adult (∼7 years old), male rhesus macaques who completed a variant of the visual reinforcement learning task previously published in ^22^ with simultaneous pupillometry recording. The pupil data were acquired in non-contiguous trials lasting several seconds (∼2-10 seconds), with each data segment corresponding to a single task trial. One study session was selected per monkey, each including more than 1500 trials of the learning task. Only trials with pupillometry recordings lasting longer than 5 seconds were selected for each monkey. This criterion resulted in the inclusion of 184 and 202 trials for each monkey, respectively, or approximately 25 minutes of pupil data per monkey.

Head-fixed pupillometry was acquired using the MATLAB-based (MathWorks, Inc.) Monkeylogic toolbox (Version 2.2; https://monkeylogic.nimh.nih.gov; ^23^) and Arrington Viewpoint eye-tracking system (recorded eye: left eye; sampling rate = 1000 Hz; Arrington Research, Inc.) run on a desktop computer (Dell OptiPlex 9020; Dell, Inc.). An infrared camera was positioned ∼40 centimeters from the monkey. Pupil size was acquired in arbitrary units, linked to the voltage output of the eye tracking system. The pupil data reported in the current study were not previously published.

### Target Pupil Phase Events

Pupil size fluctuations consist of four main phases: (1) dilation, (2) peak, (3) constriction, and (4) trough (Figure 1B). The *dilation* phase is when pupil size increases. The *peak* phase is when the pupil has reached a local maximum (i.e., the moment of a phase switch from a period of dilation to constriction). The *constriction* phase is when pupil size decreases. The *trough* phase is when the pupil has reached a local minimum (i.e., the moment of a phase switch from a period of constriction to dilation). During pupillary unrest – the spontaneous fluctuation of pupil size under constant environmental luminance – pupil size cycles among these four pupil phases, in part, according to neurophysiological state and other factors, including eye movements (e.g., blinks)^17^.

### Real Time Detection of Pupil Phase Events

Real time monitoring methods are available for various physiologic signals, for example, neural and cardiac potentials^24,25^. Meanwhile, there are limited real time monitoring methods for the pupil, in part, because pupil physiology presents unique challenges for developing real time methods. First, pupil size undergoes large baseline shifts over seconds, for example, linked to states of arousal and locomotion. These low-frequency drifts limit the efficacy of applying constant pupil size thresholds for detecting pupil phases. Second, pupil size trends do not conform to a consistent profile or shape (e.g., the stereotyped membrane potential change of a neuronal action potential). Third, spontaneous and evoked pupil size changes are slow – unfolding over 100s to 1000s of milliseconds. Therefore, at the sampling rates commonly used in pupil data acquisition (>20 Hz), there are small differences in pupil size between adjacent data samples. As a result, a derivative approach (i.e., pupil size sample x_1_-x_2_, x_2_-x_3_, etc.) to determine pupil phase is limited. Finally, the pupil data stream is subject to repeated artifacts and interruptions due to behaviors that occlude the pupil, including blinks, drooping eyelid (e.g., due to drowsiness), and whisking in rodents. *rtPupilPhase* – our method for detecting the phases of pupillary fluctuation in real time – overcomes these unique challenges of pupil physiology for developing real time pupillometry methods.

rtPupilPhase is a Python-based, open source software that we integrated with a fixation task (see *Fixation Task* section). The main goal of rtPupilPhase is to analyze live streamed pupil size data and detect the real time occurrence of target pupil phase events: dilation, peak, constriction, and trough (Figure 1B; see *Target Pupil Phase Events* section). As a control condition, rtPupilPhase also selects pupil phase-independent or *random* events. Here, we use the random events to measure the performance of rtPupilPhase. Future applications of rtPupilPhase can suppress the selection of random events or implement these events as a pupil phase-independent control or baseline condition.

The rtPupilPhase method is summarized in four main stages (Figure 1C; see Table 1 for a list of the method parameters):

#### Stage 1

Monocular, pupil size data is live streamed (see *Human Pupillometry Acquisition* section) and stored in brief, continuous intervals of pupil size data called a *pupil sample*. The pupil sample is filled with sequential pupil size data points until it comprises a total of 0.1 seconds of pupil size data (i.e., ∼6 pupil size data points at ∼60 Hz sampling rate). For the human pupillometry acquisition system, blinks, and other interruptions in the pupil data stream (e.g., lost eye tracking due to head movement or drooping eyelid) results in a pupil size value of 0. Any pupil size value in the pupil sample equal to 0 is replaced with the designation not-a-number (NaN).

#### Stage 2

The pupil sample is simultaneously added to a *search window* and *baseline window*. Subsequently, the pupil sample is cleared, and a new pupil sample is built (see Stage 1). Thus, overtime, the search and baseline windows grow, comprising continuous pupil size data that are sequentially added in increments of pupil sample (i.e., 0.1 seconds). The search window is analyzed for pupil phase events (see Stage 3). The baseline window is used to recurrently update the pupil size and pupil size derivative thresholds that determine the occurrence of pupil phase events within the search window (see Stage 4). If one or more samples in the current search window includes a NaN, the search window is cleared, and a new search window built. The current search window is also reset if the *previous* search window contained at least one NaN sample. With this approach, rtPupilPhase limits the influence of blinks and other common human pupillometry recording artifacts in the real time detection of pupil phase events by only considering pupil data intervals absent these artifactual occurrences.

#### Stage 3

When the search window comprises a minimum of two pupil samples (i.e., at least two cycles of Stages 1 and 2 have iterated without resetting the search window), the search window data is demeaned (i.e., the mean of all the pupil size data in the search window is subtracted from all search window samples). Next, the demeaned search window pupil size data is fit with a quadratic model (x-coordinates = 0 to the number of samples in the search window minus one; NumPy function *polyfit*; https://numpy.org). The resulting fitted trace is a model of the pupil size data in the search window. The final pupil size value from the fitted model timecourse, corresponding with the final sample in the search window, is recorded for later use to guide determining the pupil phase (see Stage 4). Stages 1 through 3 iterate until a target pupil phase event is detected (see Stage 4) or if the search window exceeds 5 seconds of cumulative pupil size data. In either case, the search window is emptied and built anew from subsequent pupil samples (see Stages 1 and 2).

#### Stage 4

When there is at least one previous fitted model (i.e., Stage 3 has iterated two or more times), the previous and current final pupil size values from the fitted model timecourses are compared to determine the occurrence of a target pupil phase event. A peak event is detected when the final pupil size value of the current search window is *less than* the final pupil size value from the previous search window (final value of search window_i_ < final value of search window_i-1_) *and* the final pupil size value from the current search window is *greater than* the pupil size 75-percentile for all peak events detected in the previous baseline window (see details below). A trough event is found when the final pupil size value of the current search window is *greater than* the final pupil size value from the previous search window (final value of search window_i_ > final value of search window_i-1_) *and* the final pupil size value from the current search window is *less than* the pupil size 25-percentile for all trough events found in the previous baseline window (see details below). The dilation and constriction events are detected when the difference between the final pupil size values from the fitted model timecourses of the current and previous search windows (final value of search window_i_ – final value of search window_i-1_) is *greater than* the pupil size derivative 99-percentile or is *less than* the pupil size derivative 1-percentile, respectively, the derivative percentile thresholds evaluated across all valid pupil size samples in the previous baseline window (see details below).

In the initial iterations of Stage 4, the pupil size and pupil size derivative thresholds (Table 1) are set to default constants (peak pupil size threshold = 0; trough pupil size threshold = 0; dilation pupil size derivative threshold = 50; constriction pupil size derivative threshold = -50). Subsequently, these thresholds are updated when the baseline window is filled – approximately every 5 seconds. At least half of the baseline window samples must be valid pupil size values (i.e., non-NaNs) to accept the current baseline window for analysis to update the pupil size and pupil size derivative thresholds. If the baseline window is rejected, the threshold values remain the same and the baseline window is cleared and filled again (see Stage 2). However, when a valid baseline window is confirmed, the baseline window pupil size data is demeaned, and all NaN pupil size samples are removed from the baseline window. The pupil size threshold is determined by finding all peak and trough events (SciPy function *find_peaks*; https://scipy.org) in the demeaned baseline window and then calculating the pupil size 75-percentile among peaks and pupil size 25-percentile of among troughs. Prior to finding troughs in the baseline window, the pupil size data are inverted (i.e., multiplied by negative 1), so that finding peaks in the inverted data identifies troughs in the original, uninverted dataset. The pupil size at each trough event is calculated with the original data. The pupil derivative thresholds are calculated by finding the derivative across all valid pupil size samples in the baseline window (NumPy function *diff*; https://numpy.org) and then calculating the 99 and 1-percentile of all derivative values.

When a pupil phase event is detected, the search window is cleared, and the procedure begins anew from Stage 1. An inter-event interval (IEI) of 3 seconds is set so that rtPupilPhase will not log an *accepted* pupil phase event unless the IEI has been exceeded. Events detected within the IEI are logged but designated as *not* accepted. When the IEI is exceeded for any detected pupil phase event, the search window is *not* cleared, instead additional pupil samples are added to the search window until a pupil event is detected that exceeds the IEI or the maximum length of the search window (5 seconds) is exceeded, in either case, the search window is reset. We implemented the IEI in the current demonstration of rtPupilPhase to limit the number of events analyzed to manage the computational load and so that the data samples contained within the pupil size, blink, and saccade fraction epochs are non-overlapping with any other event epoch. Future application of rtPupilPhase can modify the IEI duration or eliminate it altogether (i.e., set the IEI to 0 seconds).

### Human Simulated Real Time Detection of Pupil Phase Events

Simulated rtPupilPhase or *sim-rtPupilPhase* is a MATLAB-based (MathWorks, Inc.), open source software designed to simulate the rtPupilPhase method in a live recording using previously acquired pupil data. sim-rtPupilPhase was developed as a proof of concept to assess the performance of rtPupilPhase on pupil data from non-human animals. In future applications, sim-rtPupilPhase can serve as a tool for optimizing the rtPupilPhase parameters to achieve specific experimental goals and without collecting additional data. In summary, sim-rtPupilPhase loads previously collected pupil data and streams the data through the rtPupilPhase method (see *Real Time Detection of Pupil Phase Events* section; Figure 1C). The pupil data was modeled (see Stage 3 of the rtPupilPhase method) with the MATLAB function *fit* (x-coordinates = 1 to the number of samples in the search window; 0 to the number of samples in the search window minus 1 was also tested and achieved similar results [data not shown]; MathWorks, Inc.). To directly compare the performance between rtPupilPhase and sim-rtPupilPhase on the human pupil data, the pupil size data were down sampled from its offline sampling rate of 1000 to 60 Hz (i.e., the approximate online sampling rate of the human pupillometry recordings; see *Human Pupillometry Acquisition* section). Otherwise, the sim-rtPupilPhase parameters (e.g., the pupil size and pupil size derivative thresholds) were set to exactly as those used in the rtPupilPhase live study (Table 1).

### Mouse Simulated Real Time Detection of Pupil Phase Events

All pupil size samples concurrent with a locomotion event or with a value less than 0 (e.g., indicative of blinking or whisking) were replaced in the pupil data with NaN (see *Mouse Pupil Data* section). Otherwise, all sim-rtPupilPhase parameters and procedures were exactly as those used in the human sim-rtPupilPhase testing (Table 1; see *Human Simulated Real Time Detection of Pupil Phase Events* section).

### Monkey Simulated Real Time Detection of Pupil Phase Events

To accommodate the brief pupil data segments (<10 seconds) acquired in monkeys corresponding with task trials (see *Monkey Pupil Data* section), several adjustments were made to the sim-rtPupilPhase parameters (Table 1). First, the baseline window duration was adjusted from 5 seconds, as used in humans and mice, to 0.5 seconds. In addition, where in humans and mice, all event types were considered together to determine if the IEI threshold was breached, in the monkey dataset, each pupil phase event was considered relative to its own event type to determine if the IEI was exceeded (i.e., events of different type could fall within the IEI). This adjustment was made to increase the number of accepted pupil phase events detected in the monkey pupil data. Another adaptation made in the monkey sim-rtPupilPhase procedure was the initial removal of artifactual periods within each trial using the human pupil size preprocessing procedure (see *Monkey Pupil Data* and *Human Pupil Size, Blink, and Saccade Fraction Epoch Extraction* sections).

Preprocessing was necessary because the monkey pupil data contained numerous artifacts, including those linked to eye movements and task stimuli. Otherwise, all sim-rtPupilPhase parameters and procedures were exactly as those used in human sim-rtPupilPhase (Table 1; see *Human Simulated Real Time Detection of Pupil Phase Events* section).

## Statistical Analysis

### Human Pupil Size, Blink, and Saccade Fraction Epoch Extraction

Epoch extraction involved selecting segments of pupil size, blink, and saccade occurrence data centered on the pupil phase and random event times that were detected by rtPupilPhase and sim-rtPupilPhase. Prior to extracting the pupil size epochs, the pupil data were preprocessed for each participant, including the removal of blink and other artifactual events (e.g., drooping eyelid) that result in blank periods in the pupil size data stream (*stublinks.m* available at http://www.pitt.edu/~gsiegle)^26^. The preprocessing procedure also outputs a binary blink vector of the same length as the inputted pupil size data – blink data samples indicated with 1 and non-blink samples with 0. Similarly, a saccade binary vector was made using an established saccade detection method^27,28^ – saccade data samples indicated with 1 and non-saccade samples with 0. Note that the saccade binary vector was inclusive of microsaccades. Therefore, each participant contributed a preprocessed pupil size, blink binary, and saccade binary vector that extended the entire pupillometry recording session.

The preprocessed pupil size, blink binary, and saccade binary data were cut into 5001-millisecond epochs centered on each pupil phase and random event (i.e., 2.5 seconds before and after the event time) detected with either rtPupilPhase or sim-rtPupilPhase. The epoch was not extracted if 2.5 seconds before or after the event time exceeded the length of the pupillometry recording. The extracted pupil size epochs were demeaned (i.e., the mean pupil size across the entire epoch was subtracted from all epoch samples). After extracting all epochs, the mean across epochs was calculated within participant and events detected with rtPupilPhase and sim-rtPupilPhase, so that each participant contributed a rtPupilPhase and sim-rtPupilPhase mean pupil size, blink, and saccade fraction timecourse per pupil phase and random event. To interrogate the sensitivity of rtPupilPhase to eye movements (e.g., blinks and saccades), a subset of pupil size epochs with neither blinks nor saccades in the 0.5 seconds prior to the event time were selected and averaged separately within participant and rtPupilPhase and sim-rtPupilPhase methods (Supplementary Figure 6A, B). The participant-level saccade timecourses were smoothed using the MATLAB *smooth* function (window size = 0.1 seconds; MathWorks, Inc).

Subsequent group-level analyses involved finding the mean across participant pupil size, blink, and saccade fraction timecourses within event type and rtPupilPhase and sim-rtPupilPhase detection methods (Supplementary Figure 4A, B; Supplementary Figure 6A, B). An alternative group-level analysis first calculated the z-score across participant timecourses and then calculated the mean of the z-scored timecourses across participants within event and method type (Figure 2A, B; Supplementary Figure 3A, B). The standard error of the mean was calculated across participants. For visualization of the pupil size, blink, and saccade fraction timecourses, only the 1.5 seconds before and after the event were displayed (Figure 2A, B; Supplementary Figures 2B; 3A, B; 4A, B; 6).

### Mouse Pupil Size Epoch Extraction

The mouse pupil data were cut into 5001-millisecond epochs (101 samples at ∼20 Hz sampling rate; see *Mouse Pupil Data* section) centered on each pupil phase and random event detected with sim-rtPupilPhase (i.e., 2.5 seconds before and after the event time). The epoch was not extracted if 2.5 seconds before or after the event time exceeded the length of the pupillometry recording. Next, the pupil size epochs were demeaned (i.e., the mean pupil size across the entire epoch was subtracted from all epoch samples). After extracting all epochs, the mean across epochs was calculated within mouse and event type, so that each mouse contributed a single mean pupil size timecourse for each event. Group-level epoch analyses involved finding the mean across mouse pupil size timecourses within event type (Supplementary Figure 4C). An alternative group-level analysis first calculated the z-score across mouse pupil size timecourses and then calculated the mean of the z-scored timecourses across mice (Figure 2C; Supplementary Figure 3C). The standard error of the mean was calculated across mice. For visualization of the pupil size timecourses, only the 1.5 seconds before and after the event were displayed (Figure 2C; Supplementary Figure 3C; Supplementary Figure 4C).

### Monkey Pupil Size Epoch Extraction

Prior to extracting the monkey pupil size epoch timecourses, the pupil data were preprocessed using the same method applied on the human pupil data (see *Human Pupil Size, Blink, and Saccade Fraction Epoch Extraction* section). The monkey pupil data were cut into 1001-millisecond epochs centered on the pupil phase and random events (i.e., 0.5 seconds before and after the event time) that were detected by sim-rtPupilPhase (see *Monkey Simulated Real Time Detection of Pupil Phase Events* section). The epoch duration was reduced for the monkey pupil data relative to the 5001-millisecond epochs extracted from the human and mouse datasets because the monkey data were acquired in short segments (see *Monkey Pupil Data* section). Therefore, cutting longer epochs exceeded the bounds of the pupil data acquired in each data segment. Accordingly, epochs were not extracted if 0.5 seconds before or after the event time exceeded the length of the pupillometry recording. Next, the pupil size epochs were demeaned (i.e., the mean pupil size across the entire epoch was subtracted from all epoch samples). Unique to the monkey dataset, the demeaned epochs were also detrended using the MATLAB *detrend* function (MathWorks, Inc.) to correct for a low-frequency drift in pupil size related to the task the monkeys were engaged. The mean across all detrended epochs within monkey and pupil phase and random events was calculated, so that each monkey contributed a single mean pupil size timecourse for each event. Group-level epoch analyses involved finding the mean across all monkey-mean pupil size epoch timecourses within event type (Supplementary Figure 4D). An alternative group-level analysis first calculated the z-score across monkey epoch timecourses and then calculated the mean of the z-scored epochs across monkeys for each pupil phase and random event (Figure 2D; Supplementary Figure 3D). The standard error of the mean was calculated across monkeys.

### Cluster-Based Permutation Testing

Human pupil size timecourses for the pupil phase events were statistically tested against the random event pupil size timecourse with cluster-based permutation testing (number of permutations = 250; *p* < 0.05). The permutation analysis method employed in this study is a modified version of the method shared in the Mass Univariate ERP Toolbox (https://openwetware.org/wiki/Mass_Univariate_ERP_Toolbox)^29^. This approach was also used in a previous experiment to statistically analyze pupil size timecourses^5^. In summary, the cluster-based permutation testing method involves creating positive and negative null statistical distributions by participant-level permutation between the compared event types (e.g., peak versus random event).

Subsequently, a one-sample *t*-test was performed on the non-permuted data, identifying statistically significant positive and negative clusters based on the null distributions. Clusters were defined by temporal adjacency (i.e., statistically significant data samples contiguous in time). Four cluster-based permutation analyses were evaluated each for the rtPupilPhase and sim-rtPupilPhase human pupil size timecourses: (1) dilation versus random, (2) peak versus random, (3) constriction versus random, and (4) trough versus random events (Supplementary Figure 4A, B). The mouse and monkey data were not subjected to statistical testing due to low individual animal sample sizes.

### Pupil Phase Event Detection Accuracy and Sensitivity Performance

The accuracy and sensitivity of rtPupilPhase in detecting pupil phase events in humans was calculated based on accepted events (i.e., those that exceeded the IEI; see *Real Time Detection of Pupil Phase Events* section). Event accuracy was evaluated by comparing the real time detected pupil phase event versus the corresponding *true* pupil phase. The true pupil phase was determined by post hoc analysis of the pupil data. In summary, first, the human pupil data were preprocessed, including removing blinks and artifactual samples (see *Human Pupil Size, Blink, and Saccade Fraction Epoch Extraction* section). Next, the preprocessed pupil data were smoothed using the MATLAB *smoothdata* function (Savitzky-Golay filter; window size = 0.1 seconds; MathWorks, Inc.) to eliminate high-frequency signals. The true peak and trough pupil phase events were identified using the MATLAB *findpeaks* function (MathWorks, Inc.). For trough events, prior to implementing findpeaks, the pupil size data were inverted (i.e., multiplied by negative 1).

Finally, the top 75 percent of peaks and troughs were selected based on their prominence. For two participants, the prominence percent threshold was adjusted to 90 percent because visual inspection determined that lower thresholds resulted in the selection of peak and trough events with low prominence. Finally, peak and trough event binary indices were made by setting all samples within 0.25 seconds before and after each peak and trough event to an index value of 1, while all other samples outside this range were set to 0.

The true dilation and constriction pupil phase events were found by calculating the slope at every sample of the pupil size data using the MATLAB *gradient* function (MathWorks, Inc.). A dilation event binary index was created by setting all samples from the gradient output with a positive value to 1, indicating a positive slope or dilation, and a negative value to 0, indicating a negative slope or constriction. Conversely, the constriction binary index was created by setting all the samples from the gradient output with a negative value to 1 and a positive value to 0.

The accuracy of the real time detected pupil phase events was assessed by comparing each event sample to the corresponding true pupil phase event index. An index value of 1 at the queried event sample indicated agreement between the real time and true pupil phase events. The real time detected event accuracy percentage was calculated within participant and pupil phase event type detected with rtPupilPhase by determining the total number of pupil phase events that were accurately detected (i.e., in agreement with the true pupil phase event indices) divided by the total number of detected pupil phase events. The result was multiplied by 100 to convert from rate to percentage units. The baseline or chance accuracy rate for each pupil phase event was determined by comparing the random event samples to each of the true pupil phase event indices (e.g., random event versus true dilation event index, random event versus true peak event index, etc.). Finally, to assess if the real time pupil phase event accuracy was greater than the random accuracy percentage, group-level paired *t*-tests across participants were conducted to statistically compare pupil phase and random event accuracies (Holm-Bonferroni corrected; *p* < 0.05). A total of four *t*-tests were completed: (1) dilation versus random, (2) peak versus random, (3) constriction versus random, and (4) trough versus random events (Figure 3B).

Finally, the sensitivity of the real time detected pupil phase events was assessed using the exact method as detailed above for calculating rtPupilPhase accuracy except each real time detected event sample was compared to *off-target* true pupil phase event indices (e.g., comparing the real time detected dilation event with the true peak, constriction, and trough event indices). A one-sample *t*-test (Holm-Bonferroni corrected; *p* < 0.05) tested if the accuracy of the real time detected pupil phase events (i.e., on-target performance; e.g., comparing the predicted dilation event with the true dilation event index) was greater than the off-target performance (Supplementary Figure 5). A statistically greater on-target accuracy suggests that rtPupilPhase is sensitive to each pupil phase event type. These analyses also help to assess overlap among the pupil phase events (i.e., a detected pupil phase event belonging to two or more true pupil phase event types).

### Pupil Phase Event Detection Temporal Performance

The temporal performance of rtPupilPhase is defined as the inter-event duration or the time between detected pupil phase events, independent of event type. Temporal performance is impacted by the online pupillometry sampling rate, the rtPupilPhase parameters, and behaviors that occlude the pupil (e.g., blinks). The human rtPupilPhase temporal performance was calculated for each participant by, first, combining the time indices for all detected pupil phase events, ignoring the IEI and random events (see *Real Time Detection of Pupil Phase Events* section) and sorting these events in chronological order. Also, intervals between fixation task blocks (i.e., ∼2-minute task break periods; see *Fixation Task* section) were ignored. Next, the duration of time between each event was calculated (i.e., t’ = event time_1_ minus event time_2_), independent of event type. The median t’ value was calculated for each participant and then averaged across participants to determine the estimated temporal resolution of rtPupilPhase (Figure 3B). We choose the median to statistically assess temporal performance because there were prolonged durations between detected pupil phase events (t’ > 0.5 seconds in 15.34% of inter-event durations commonly the result of blinking bouts; Figure 2B) that biased mean-based analyses. Finally, the detection frequency rate was calculated by evaluating 1 divided by the group median temporal resolution converted to units of seconds.

## Data and Code Availability

All code and data are available at https://github.com/nimh-sfim/rtPupilPhase.

## Author Contributions

S.I.K. contributed to conceptualization, methodology, software, formal analysis, investigation, data curation, visualization, supervision, project administration, and writing (original draft); V.E.G. contributed to methodology, software, formal analysis, investigation, and writing (review and editing); C.R.W. contributed to software, data curation, and writing (review and editing); J.B.T. contributed to methodology, software, and writing (review and editing); D.C.B. contributed to formal analysis, resources, and writing (review and editing); S.S. contributed to formal analysis, resources, and writing (review and editing); J.G-C. contributed to conceptualization, methodology, supervision, and writing (review and editing); P.A.B. contributed to conceptualization, methodology, supervision, funding acquisition, and writing (review and editing).

## Acknowledgements

This research was made possible by the support of the National Institute of Mental Health Intramural Research Program (ZIAMH002783 and ZIAMH002928). The study was completed in compliance with the National Institutes of Health Clinical Center protocol ID 93-M-0170 (ClinicalTrials.gov ID: NCT00001360). A special thank you to Daniel A. Handwerker for his constructive feedback on the project scripts and GitHub repository.

---

**Supplementary Figure 1.**
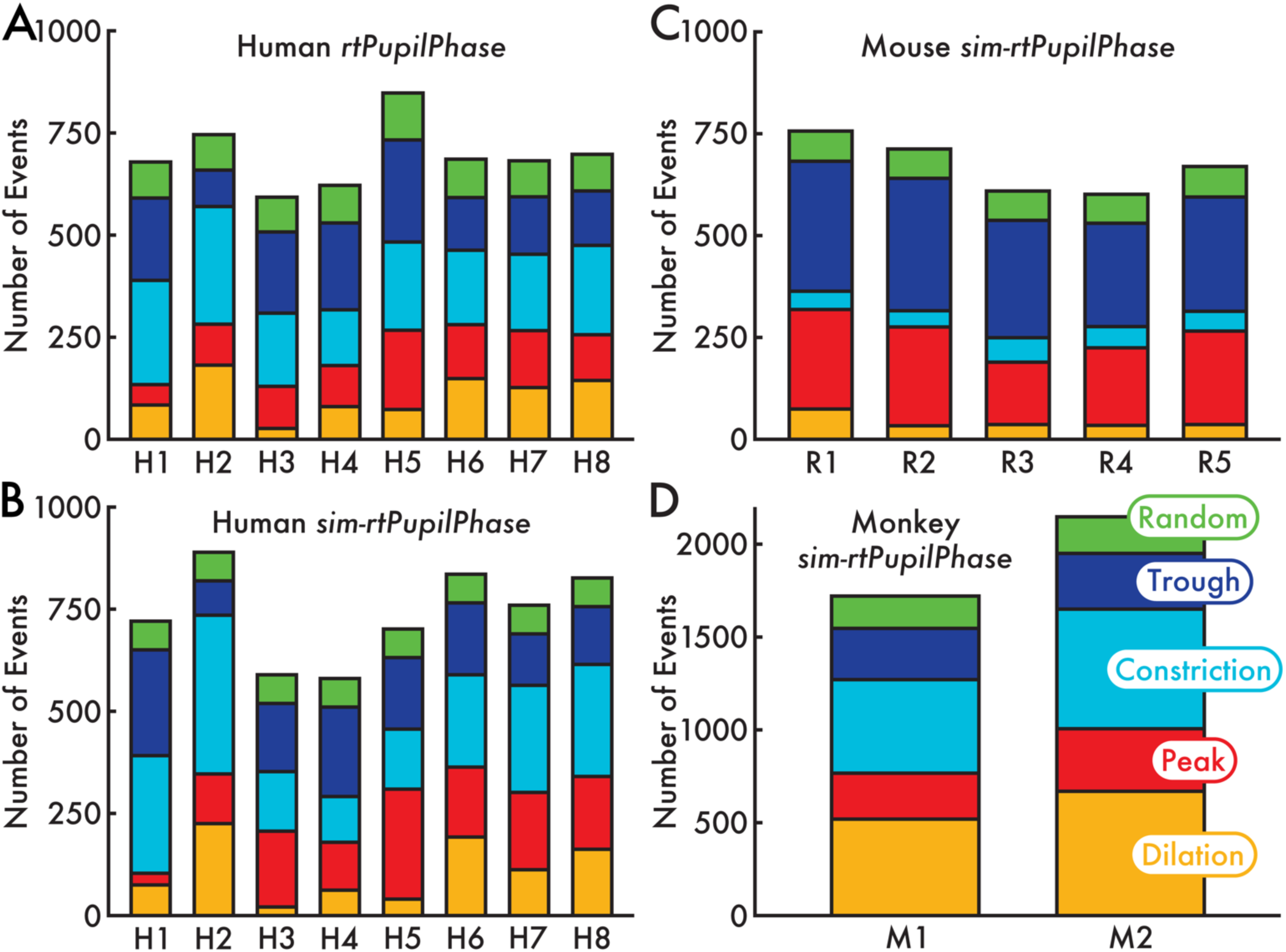
Number of pupil phase and random events detected across individual humans, mice, and monkeys. **(A)** Human (H; N = 8) rtPupilPhase detected pupil phase and random events. **(B)** Human (N = 8) sim-rtPupilPhase detected pupil phase and random events. **(C)** Mouse (R; N = 5) sim-rtPupilPhase detected pupil phase and random events. **(D)** Monkey (M; N = 2) sim-rtPupilPhase detected pupil phase and random events. The depicted events are the same as those reported in Figures 2, 3 and Supplementary Figures 2-6. Modulating the parameters of rtPupilPhase (Table 1) can adjust the number of detected events. The bar color specifies the event type.

**Supplementary Figure 2.**
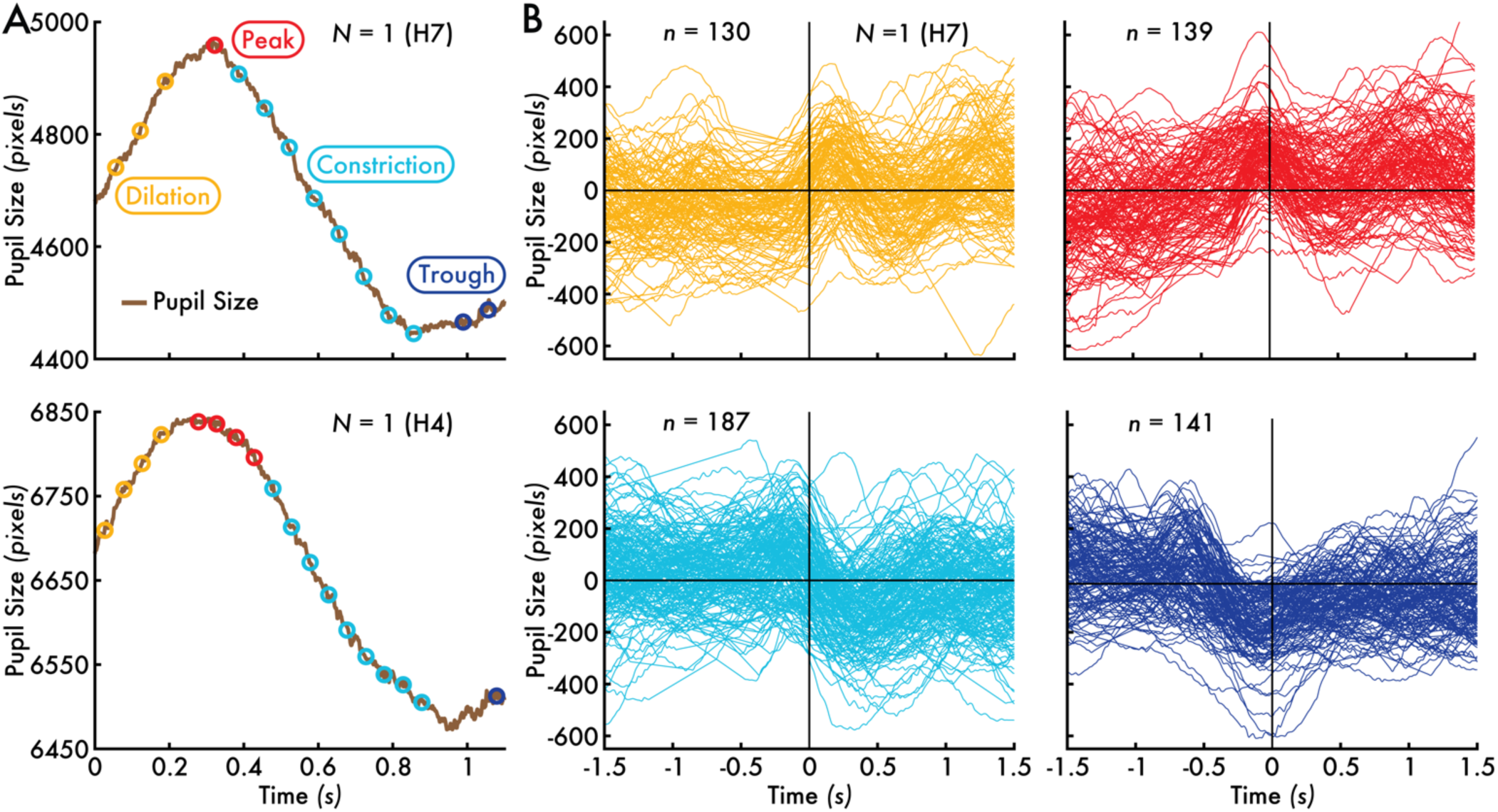
Event-level rtPupilPhase performance. **(A)** Pupil size trace interval (1.1 seconds [s]) from two human participants (H4, H7; see Supplementary Figure 1). Open circles indicate the time of real time pupil phase events detected with rtPupilPhase, ignoring the inter-event interval that determines accepted events (see *Real Time Detection of Pupil Phase Events* Methods section). Between these two sample intervals (H4, H7), there was on average 15 detected pupil phase events within 1 s, corresponding to the group median pupil phase inter-event duration of 0.067 s (Figure 3B). **(B)** Individual participant (H7) pupil size timecourses for each extracted epoch centered on the pupil phase event detection time (0 s). The open circle and trace colors specify the pupil phase event type. N = number of participants; n = number of pupil phase event epochs.

**Supplementary Figure 3.**
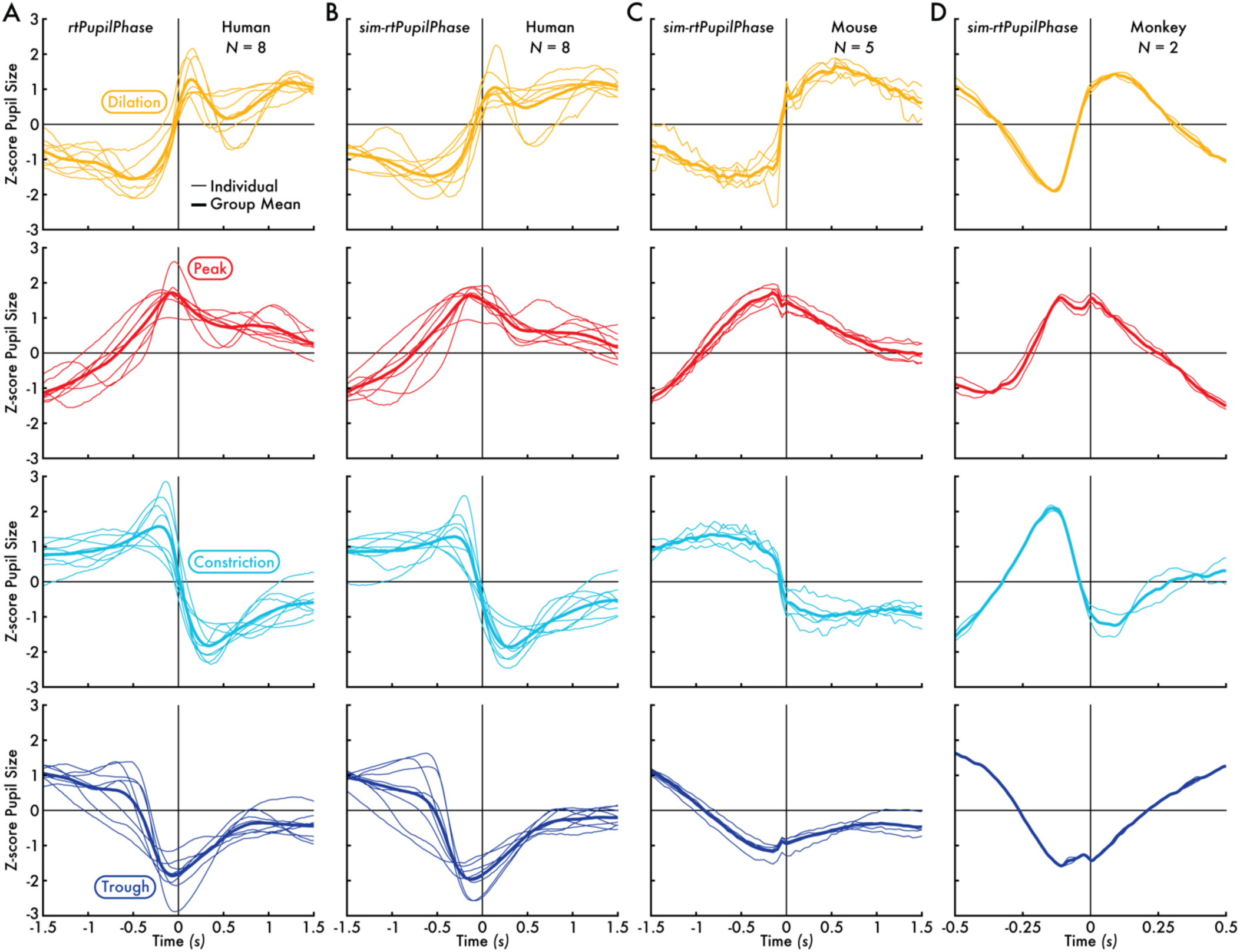
Individual human, mouse, and monkey pupil size epoch timecourses for pupil phase events detected with rtPupilPhase and sim-rtPupilPhase. **(A)** The rtPupilPhase human (N = 8) individual and group mean z-score pupil size timecourses. **(B)** The sim-rtPupilPhase human (N = 8) individual and group mean z-score pupil size timecourses. **(C)** The sim-rtPupilPhase mouse (N = 5) individual and group mean z-score pupil size timecourses. **(D)** The sim-rtPupilPhase monkey (N = 2) individual and group mean z-score pupil size timecourses. The thinner lines show the individual animal (i.e., human, mouse, and monkey) pupil size timecourses. The thicker lines show the mean pupil size timecourse across all individuals within the human, mouse, and monkey datasets (i.e., the same z-score pupil size timecourses shown in Figure 2). The pupil phase event time is 0 seconds (s). The trace color specifies the pupil phase event type.

**Supplementary Figure 4.**
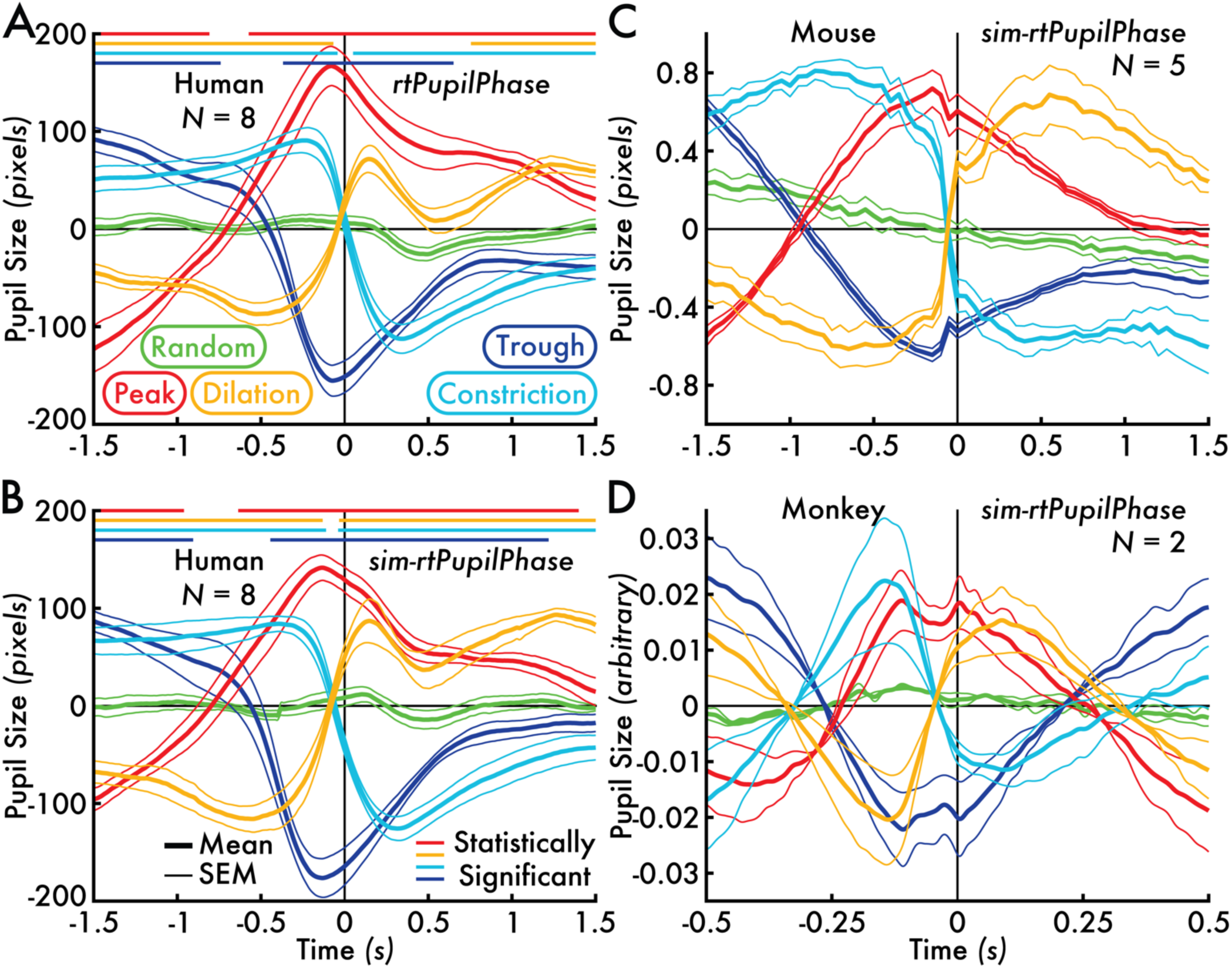
Human, mouse, and monkey pupil size timecourses centered on pupil phase events detected with rtPupilPhase and sim-rtPupilPhase in native recording units. **(A)** The rtPupilPhase human (N = 8) mean pupil size timecourses. **(B)** The sim-rtPupilPhase human (N = 8) mean pupil size timecourses. **(C)** The sim-rtPupilPhase mouse (N = 5) mean pupil size timecourses. **(D)** The sim-rtPupilPhase monkey (N = 2) mean pupil size timecourses. The group mean pupil phase event pupil size timecourse is shown with the thicker line, bounded by thinner traces that depicts the standard error of the mean (SEM). The pupil phase event time is 0 seconds (s). The trace color corresponds with the pupil phase event. The horizontal-colored lines near the top of subplots (A) and (B) depict the intervals where a statistically significant difference (cluster-based permutation testing, *p* < 0.05) was found between the pupil phase and random event (see *Cluster-Based Permutation Testing* Methods section). The statistical line color indicates the tested pair (orange = dilation versus random event; red = peak versus random event; cyan = constriction versus random event; blue = trough versus random event). Statistics were not completed on the mouse and monkey datasets due to low individual animal sample size.

**Supplementary Figure 5.**
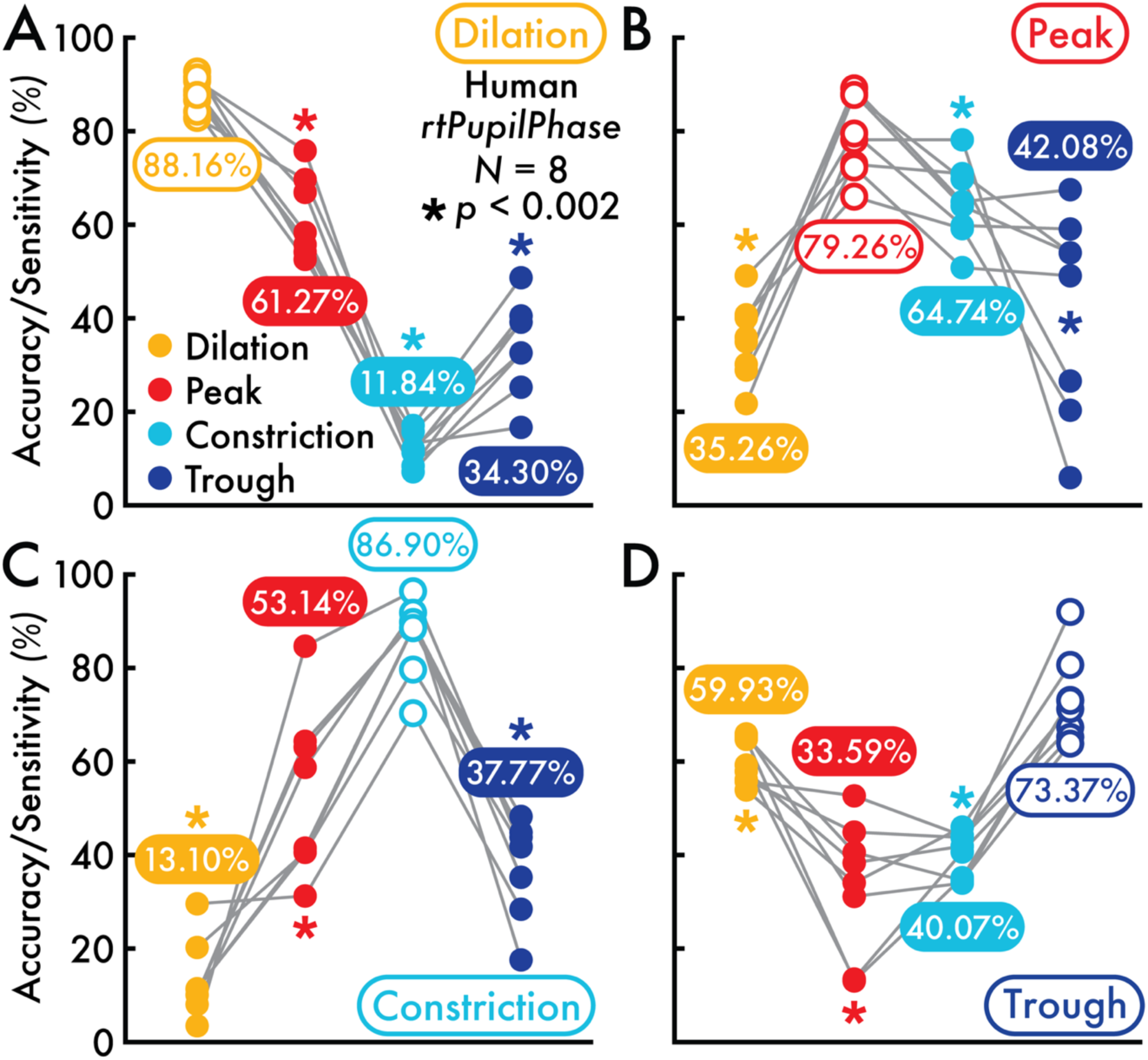
rtPupilPhase detected pupil phase event performance sensitivity. **(A)** The pupil phase accuracy percentage for each human participant (each open circle corresponds with a participant; N = 8) is shown for the rtPupilPhase real time detected dilation events versus the true dilation events (same results as reported in Figure 3A). The pupil phase sensitivity percentage for each human participant (each closed circle corresponds with a participant) is shown for the rtPupilPhase real time detected dilation events versus the true off-target pupil phase events: peak, constriction, and trough (see *Real Time Pupil Phase Event Detection Accuracy and Sensitivity Performance* Methods section). Sensitivity for detected dilation events (each circle corresponds with a participant; open circles indicate the detection accuracy performance; same results as reported in Figure 3A; N = 8) across all events. The group mean performance accuracy/sensitivity percent is shown. The dilation accuracy (88.16 percent) was statistically greater than all off-target sensitivity performances (peak = 61.27 percent; constriction = 11.84 percent; trough = 34.30 percent; Holm-Bonferroni corrected, *p* < 0.05; * *p* < 0.002). **(B)** Detected peak on-target accuracy and off-target sensitivity. On-target accuracy is significantly greater than off-target sensitivity (Holm-Bonferroni corrected, *p* < 0.05). **(C)** Detected constriction on-target accuracy and off-target sensitivity. On-target accuracy is significantly greater than off-target sensitivity (Holm-Bonferroni corrected, *p* < 0.05). **(D)**. Detected trough on-target accuracy and off-target sensitivity. On-target accuracy is significantly greater than off-target sensitivity (Holm-Bonferroni corrected, *p* < 0.05). All off-target results show that the pupil phase order corresponds with the performance sensitivity.

**Supplementary Figure 6.**
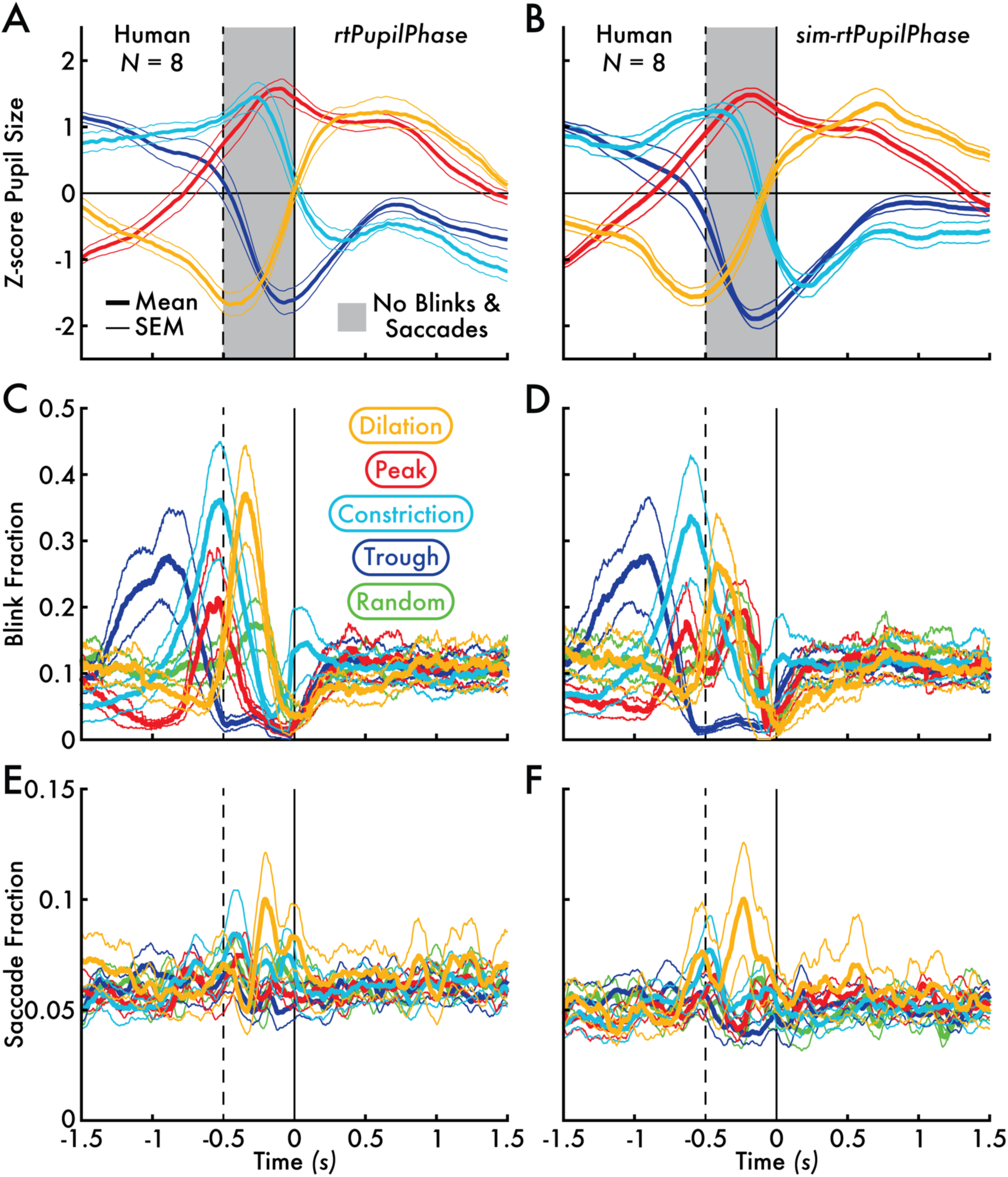
Human pupil size timecourses without blinks and saccades and blink and saccade fraction timecourses centered on pupil phase and random events detected with rtPupilPhase and sim-rtPupilPhase. **(A)** The rtPupilPhase human (N = 8) mean z-score pupil size timecourses excluding epochs with blinks or saccades between -0.5 and 0 seconds (s; exclusion interval is shown with a gray bar). **(B)** The sim-rtPupilPhase human (N = 8) mean z-score pupil phase event pupil size timecourses excluding epochs with blinks or saccades between -0.5 and 0 s (exclusion interval is shown with a gray bar). **(C)** The rtPupilPhase human mean blink fraction timecourses. **(D)** The sim-rtPupilPhase human mean blink fraction timecourses. **(E)** The rtPupilPhase mean human saccade fraction timecourses. **(F)** The sim-rtPupilPhase mean human saccade fraction timecourses. The pupil phase event time is 0 s. The vertical dashed line shown in subplots (C), (D), (E), and (F) depict the 0.5 s interval used to exclude pupil size epochs if any blink or saccade event were present (see subplots A and B). The group-level mean pupil size, blink, and saccade fraction timecourses for each pupil phase event is shown with the thicker line, bounded by thinner traces that depicts the standard error of the mean (SEM). The trace color corresponds with the pupil phase and random event type.

